# Analysis of SARS-CoV-2 ORF3a structure reveals chloride binding sites

**DOI:** 10.1101/2020.10.22.349522

**Authors:** Valeria Marquez-Miranda, Maximiliano Rojas, Yorley Duarte, Ignacio Diaz-Franulic, Miguel Holmgren, Raul E. Cachau, Fernando D. Gonzalez-Nilo

## Abstract

SARS-CoV-2 ORF3a is believed to form ion channels, which may be involved in the modulation of virus release, and has been implicated in various cellular processes like the up-regulation of fibrinogen expression in lung epithelial cells, downregulation of type 1 interferon receptor, caspase-dependent apoptosis, and increasing IFNAR1 ubiquitination. ORF3a assemblies as homotetramers, which are stabilized by residue C133. A recent cryoEM structure of a homodimeric complex of ORF3a has been released. A lower-resolution cryoEM map of the tetramer suggests two dimers form it, arranged side by side. The dimer’s cryoEM structure revealed that each protomer contains three transmembrane helices arranged in a clockwise configuration forming a six helices transmembrane domain. This domain’s potential permeation pathway has six constrictions narrowing to about 1 Å in radius, suggesting the structure solved is in a closed or inactivated state. At the cytosol end, the permeation pathway encounters a large and polar cavity formed by multiple beta strands from both protomers, which opens to the cytosolic milieu. We modeled the tetramer following the arrangement suggested by the low-resolution tetramer cryoEM map. Molecular dynamics simulations of the tetramer embedded in a membrane and solvated with 0.5 M of KCl were performed. Our simulations show the cytosolic cavity is quickly populated by both K+ and Cl-, yet with different dynamics. K+ ions moved relatively free inside the cavity without forming proper coordination sites. In contrast, Cl- ions enter the cavity, and three of them can become stably coordinated near the intracellular entrance of the potential permeation pathway by an inter-subunit network of positively charged amino acids. Consequently, the central cavity’s electrostatic potential changed from being entirely positive at the beginning of the simulation to more electronegative at the end.

## Introduction

The recent release of the SARS-CoV-2 Open Reading Frame 3a (ORF3a) dimer structure, solved by Cryo-EM [1] (PDB:6XDC), offers new opportunities for SARS-CoV antivirals design. The deletion of ORF3a reduces viral titers in animal models, suggesting ORF3a as a target for developing therapeutic agents against SARS-CoV-1 [2]; however, the exact mechanism of function of ORF3a is not well understood. ORF3a is believed to form ion channels [3] [4] [9], which may be involved in the modulation of virus release [9]. Apoptosis initiates a viral cytopathic effect in SARS-CoV-1-infected cells [5] [6] [7]. Blocking ORF3a channel activity has been reported to abolish caspase-dependent apoptosis. ORF3 is related to viral pathogenicity in porcine epidemic diarrhea virus (PEDV) [4]. Other cellular processes affected by ORF3a include the up-regulation of fibrinogen expression in lung epithelial cells, downregulation of type 1 interferon receptor, and increasing IFNAR1 ubiquitination. For a full list, see Zhang et al.[8]. The broad spectrum of responses linked to ORF3a and its distinct sequence and structure makes ORF3a a valuable target.

Initial studies [3][4][9] showed that ORF3a could form ion channels in Xenopus laevis oocytes and yeast, with an expected tetrameric assembly as inferred from biochemical data [9]. Recently, nonetheless, channel activity has been detected using purified dimeric ORF3a proteins from SARS-CoV-2 that were reconstituted in proteoliposomoes [1]. These dimers’ cryoEM high-resolution structure revealed that each protomer contains three transmembrane helices, arranging in a clockwise configuration forming a six helices transmembrane domain [1]. The potential permeation pathway of this domain is formed by TM1 and TM2 [1], consistent with functional observations [2][3]. This pathway has six constrictions narrowing to about 1 Å in radius, suggesting the structure solved is in a closed or an inactivated state. The potential permeation pathway contains a polar cavity within the inner half of the transmembrane region. The cytosol end of the structure forms a cytosolic domain composed of multiple beta strands from both protomers. Here we used this dimeric structure to perform full atom molecular dynamic simulations and electrostatic potential calculations to ask questions concerning the dimers’ stability and whether ions could be populating specific regions of the channel. We showed that the dimer core is strongly stabilized by more than 100 methyl-methyl interactions, consistent with a non-conductive conformation. Further, we evidenced that the inner polar cavity has physicochemical properties that are better suited for hosting negatively charged species instead of cations.

### Modeling of the ORF3a channel pore

Since biochemical data suggest that residue C133 is essential for tetramerization [9], we opted to establish an initial system composed of two dimers with high-resolution structures covalently joint by two C133 (Figure 1A). This tetrameric complex is compatible with the low-resolution cryoEM structure obtained from purified tetramer forms [1]. The tetramer was embedded in a membrane solvated in a water box with 0.5 M KCl. Five hundred nanoseconds molecular dynamics simulation of this system was carried out. For simulation details, see Methods. All results described below were observed in both dimers of the tetrameric system.

**Figure 1:**
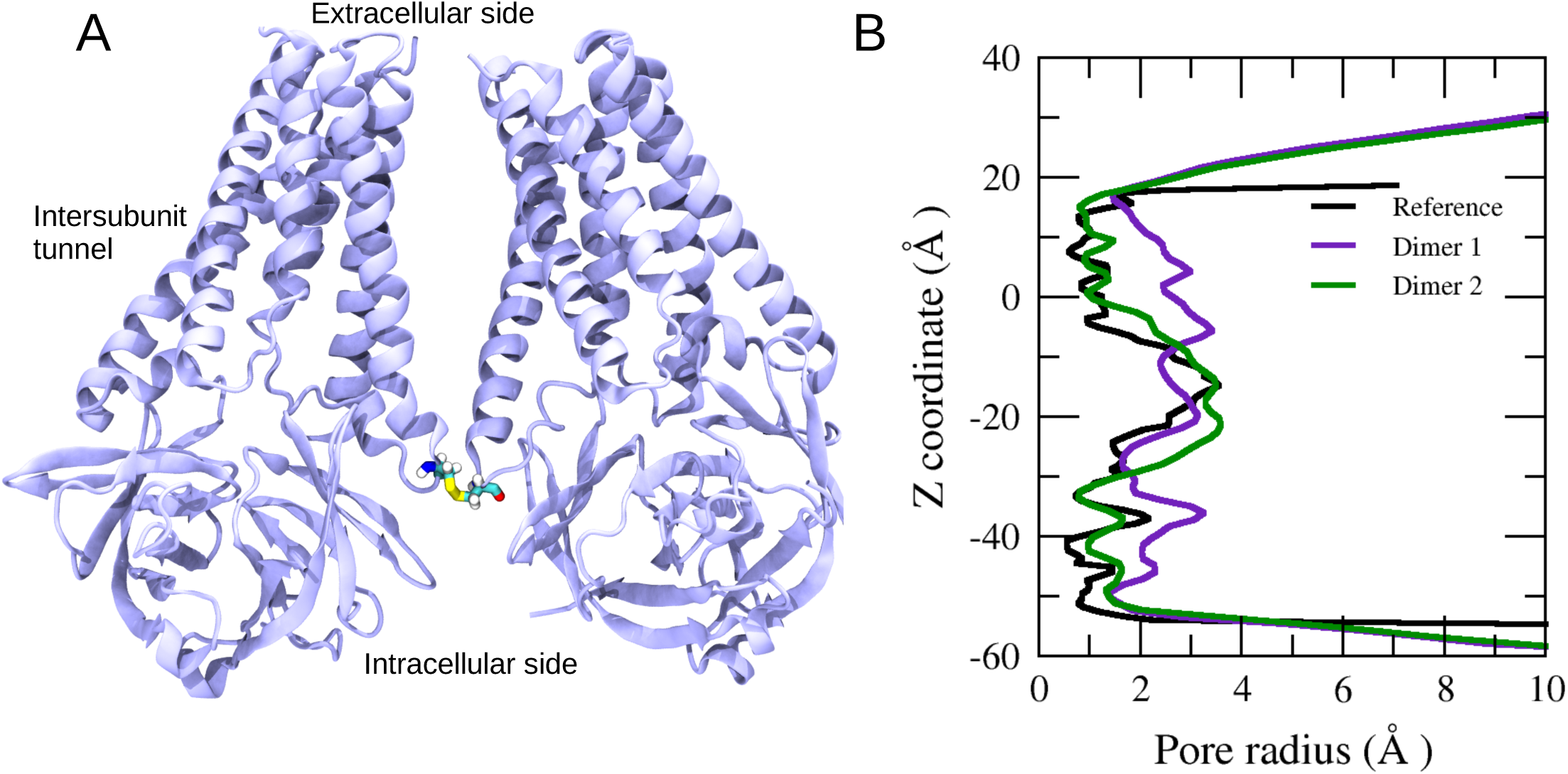
Simulation system. A) Initial ORF3a structure composed of two dimers obtained from 6XDC crystal structure, covalently joint by a disulfide bond formed by two C133 residues. B) Pore radius profile for the reference structure PDB ID: 6XDC, and for both dimers in the tetramer as obtained after 500 ns of molecular dynamics simulations. Profiles of both dimers evidence slight changes in the pore size, especially in the central polar cavity, where up to three chloride ions remain in this region (∼ z = -10 Å), widening the pore.

No significant changes in the transmembrane pore dimensions were observed during the simulation, with the pore radius widening by about 1 Å around the extracellular and intracellular side (Figure 1B). The pore dimensions stability can be explained by the strong vdW interactions dominating the interactions in the transmembrane region’s narrowest section, formed by the residues from 158 to 232. Free energy of binding between two monomers of a single dimer obtained using MM-GBSA analysis over the last 100 frames of MD indicates the vdW contribution is five times larger than that of the electrostatics (see Methods). The large vdW contribution originates in the more than 138 methyl-methyl interactions between the two monomers (chain A and chain B), resulting in a tightly packed core interaction that cannot be breached unaided. Not surprisingly, no ions were observed to cross the potential permeation pathway.

### Basic residues in the water-membrane interface are critical in the entry of chloride ions to the central polar cavity of ORF3a

In addition to the outer half of the potential permeation pathway, there are three possible access routes to the central polar cavity, so-called tunnels [1]. The upper tunnel connects the polar cavity to the lipid bilayer. Through the inter-subunit and the inner tunnels, the polar cavity opens to the bulk solution at the water-membrane interface and directly into the cytosol, respectively. The inter-subunit tunnel, located between z=-10 and z=-20 (with z measuring the vertical displacement from the membrane midpoint when the model central axis is aligned along the Z coordinate), has a high density of positively charged residues (Lys66, Lys67, Arg68), which may negatively affect cations binding while favoring negatively charged species. Indeed, our simulations show that this tunnel is the main entryway for Cl- ions into the central polar cavity. Within the first 100 ns of the simulation, three Cl- ions entered the central polar cavity and remained for the rest of the simulation. These three Cl- ions are constantly alternating their position within the polar cavity. Figure 2A shows a static view of the Cl- ions inside the central polar cavity and the amino acids responsible for their retention in this region. The presence of anions in this region makes the polar cavity slightly wider than the experimental structure dimensions. Still, it does not allow ions translocation through a potential transmembrane permeation pathway. The central polar cavity shows the highest occupancy of Cl- ions (Fig. 2B). The average density of the molecular species in the atomic system is depicted in Supp. Material Figure S1.

**Figure 2:**
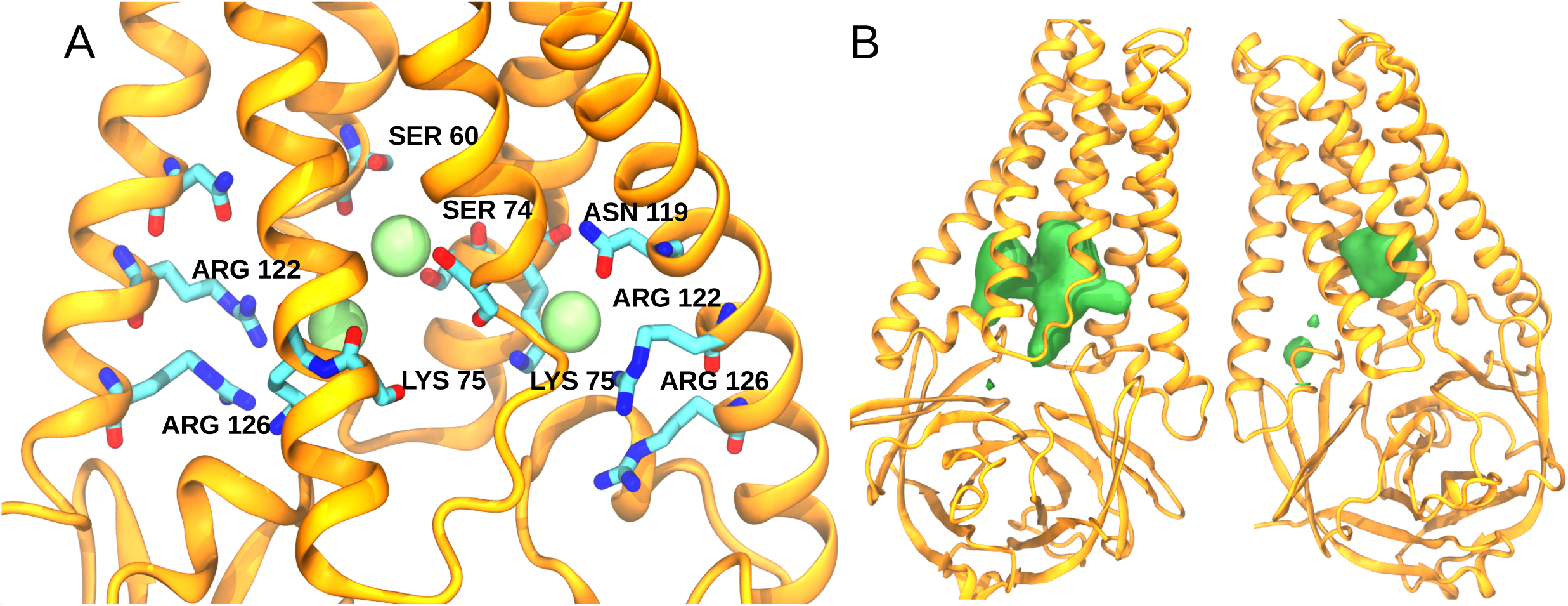
Regions of chloride ions occupancy. A) A close up of the central polar cavity shows three Cl- ions coordinated by residues of the two monomers (Arg122, Arg126, Lys75, Ser74, Ser60, and Asn119). B) Highest occupancy regions of Cl- ions, depicted in green, in the tetrameric model of ORF3a taken as an average of the 500 ns of molecular simulation.

### Potassium ions accumulate on the surface of the cytosolic domain of ORF3a

Regions with high K+ ions occupancy were assessed using the Volmap Tool in VMD[10]. The cytosolic domain, composed mainly of beta-sheets, acts as a repository of K+ ions, accumulating through the surface exposed to the cytosol (Figure 3A). In contrast to Cl- ions, no high occupancy sites for K+ were found inside the central polar cavity. We observed that most of the cytosolic domain regions where K+ ions accumulate do not form proper coordination sites, which means that K+ does not remain in those regions for a long time. There was only one K+ that remained stable at the cytosolic domain’s surface, coordinated by E181, D183, S180, and D173, along with two water molecules (Figure 3B). Interestingly, residues E171 and D173 are relevant for the potassium channel activity of SARS-CoV-1 ORF3a [11], with alanine mutations decreasing conductance when compared to the WT channel.

**Figure 3:**
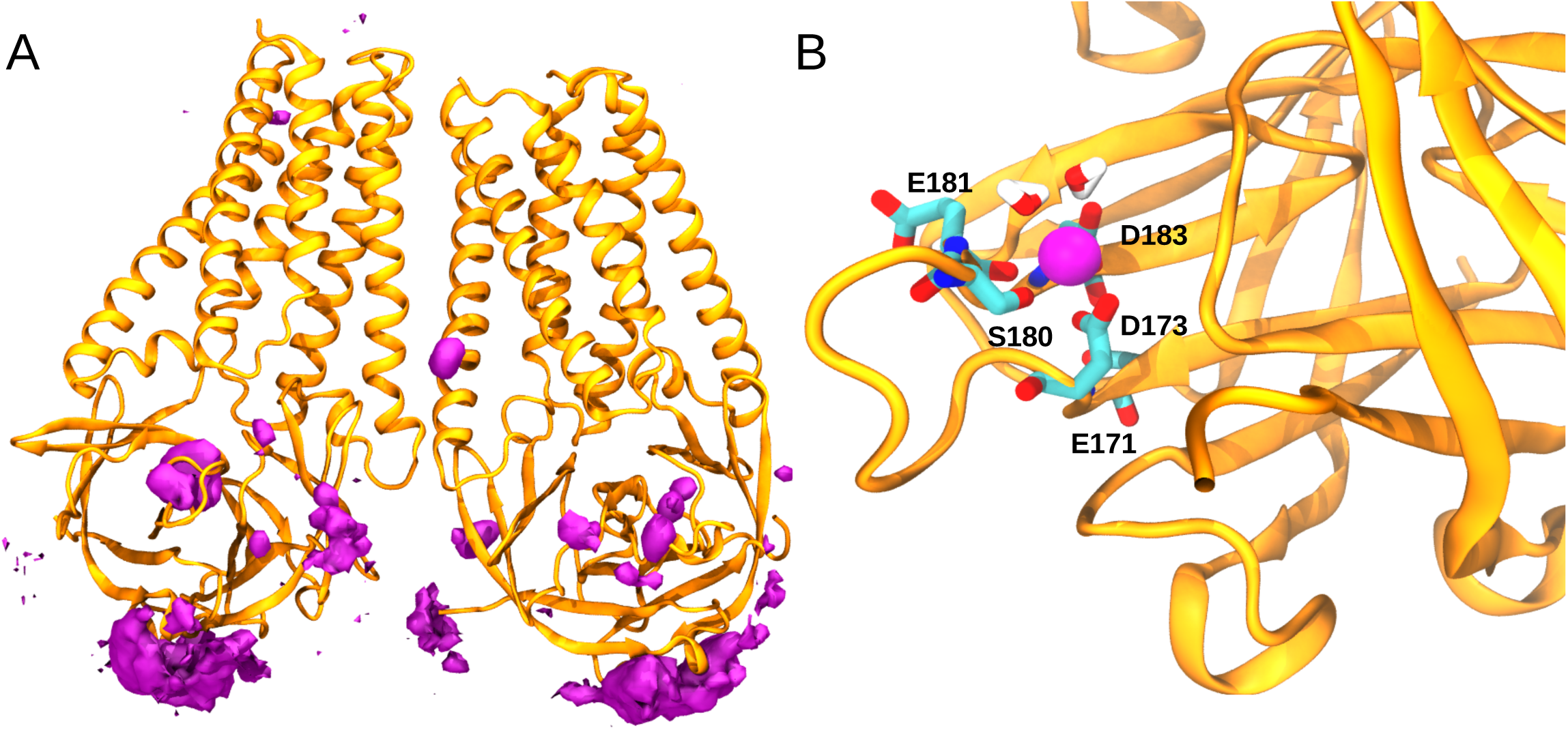
Regions of potassium ions occupancy. A) Highest occupancy regions of K+ ions, depicted in magenta, in the tetrameric model of ORF3a taken as an average of the 500 ns of molecular simulation. B) A high occupancy site (100% occupancy during the last 150 ns of MD) for a single K+ is located in the cytosolic domain. Residues E181, D183, S180, and D173, along with two alternating water molecules, are coordinating a single K+ (in magenta).

### Ion occupancies modify the electrostatic potential of the ORF3a channel

To assess the impact of the ion occupancies described above, we obtained the electrostatic potential maps for the ORF3a channel for the initial configuration, and the last frame, at the end of a trajectory of 500 ns of the molecular dynamics simulations, by employing the Poison-Boltzmann approach implemented in the APBS package [12]. This analysis shows that the entry of Cl- ions through the inter-subunit tunnel into the central polar cavity and the accumulation of K+ ions at the cytosolic domain’s surface changed the channel’s electrostatic profile. The central cavity’s electrostatic potential appears to be entirely positive in the first frame, changing dramatically after the Cl- ions occupied the central polar cavity, becoming more electronegative (Figure 4A and 4B). Interestingly, this change propagated through the transmembrane domain’s low dielectric field to the extracellular side, promoting a less positive electrostatic potential of the transmembrane region. The accumulation of K+ ions at the cytoplasmic domain’s surface also produced substantial changes in the electrostatic potential of the channel, particularly at the intracellular end between the two dimers (Figure 4A and 4B).

**Figure 4:**
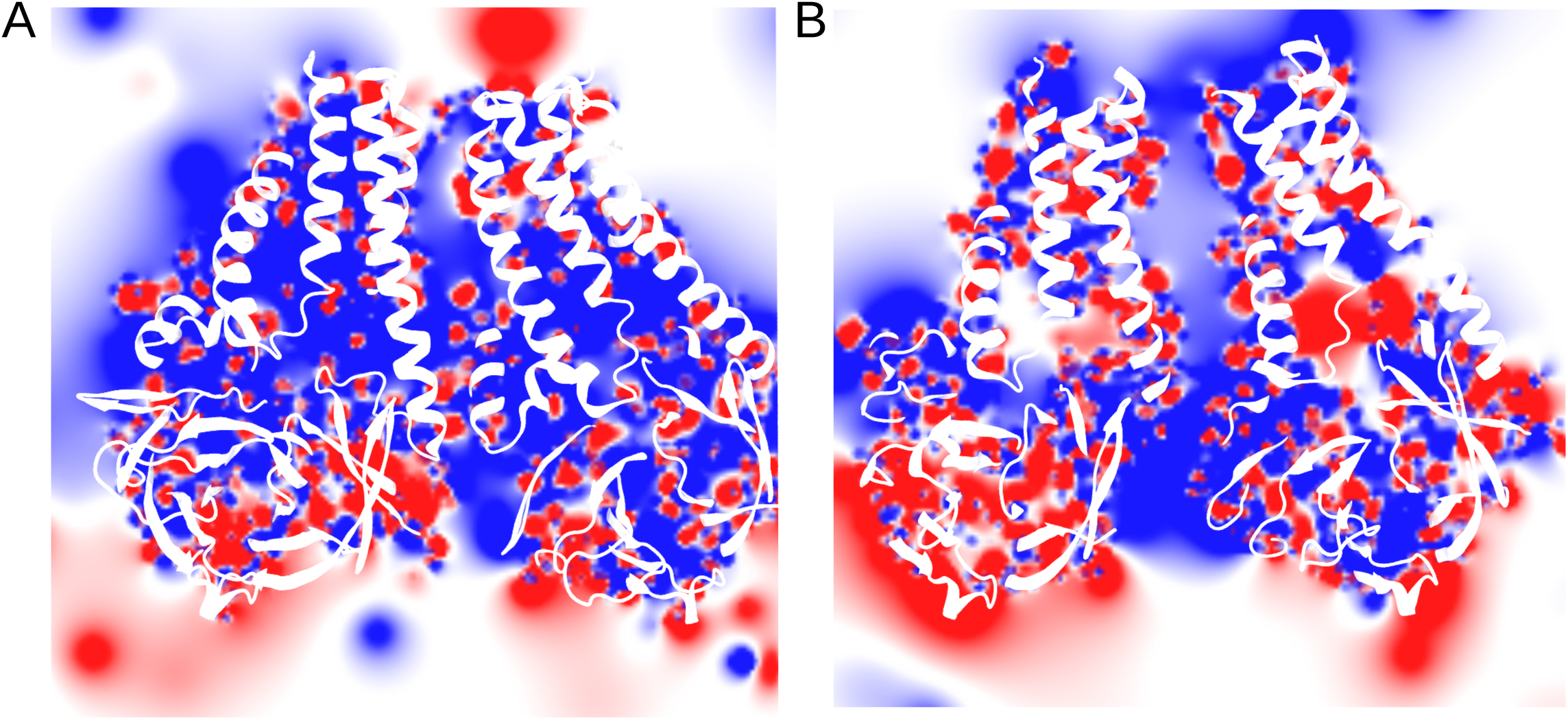
Electrostatic potential maps were obtained from the ORF3a tetramer model at the first frame of MD (A) and the last frame (B). The maps show the change in the electrostatic potential in the intersubunit tunnel, which changes from electropositive to electronegative due to chloride ions, and in the interface between beta-sheets from both ORF3a dimers, where it becomes electropositive due to the accumulation of potassium ions.

The AFAL server [13] was used to correlate our observations with previously characterized cation and anion binding sites. The AFAL server analyzes and catalogs the propensity of amino acid residues to interact with a specific ligand, based on the information available in the structures deposited in the Protein Data Bank. The residues surrounding Cl- ions with the highest probability are ARG > ASN > LYS > SER > THR, in agreement with the residues observed at the entrance of the inter-subunit tunnel and those in the central polar cavity, specifically the residues Lys66, Lys67, Arg68, Arg122, Arg126, Lys75, Ser60, Ser74, and Asn119. The AFAL profile for K+ ions showed that the higher amino acid propensity near the ion is ASP > SER > THR > GLU, consistent with the amino acids that stabilized the single K+ at the surface of the cytoplasmic domain. This information supports our observation that the ORF3a central region is not well suited to interact with positively charged species.

## Conclusions

Functional studies demonstrate that ORF3a has properties of cation-selective channels [3][4][9], an activity that has been linked to apoptosis in host cells [14]. The recent cryoEM high-resolution structure of a dimer form of ORF3a from SARS-CoV-2 and the expected tetrameric complex arrangement are unusual [1]. Here we presented an initial analysis of the SARS-CoV-2 ORF3a structure, explored with molecular simulations methods. We found that the central polar cavity has high-affinity sites for chloride ions. In contrast, no high-affinity regions for potassium ions were found in the potential permeation pathway. We only saw a high occupancy site for a single K+ in the cytosolic domain, close to the residues E171 and D173. Multiple K+ ions tend to concentrate intermittently in the cytosolic domain’s cytoplasmic surface, acting as a repository of cations.

ORF3a from SARS-CoV-1 (Uniprot:P59632) and SARS-CoV2 (Uniprot: P0DTC3) shares 85% of sequence similarity (as obtained using EMBOSS Needle [15]), yet all the key residues in the chloride anion binding sites are conserved. This observation is further supported by our analysis of distal sequences, mostly from bat viruses, where the key chloride coordinating residues are conserved (not shown). These observations indicate that the inter-subunit tunnel and the central polar cavity have evolved to coordinate negative ions. Furthermore, the negative ions are stabilized in the central polar cavity located in the inner half of the potential permeation pathway, precisely in the middle of the transmembrane protein core. Thereby, the low dielectric transmembrane region may contribute to propagating the electrostatic potential, which could help regulate the flow of cations through an alternative pore formed by a rearrangement of ORF3a monomers.

It is difficult to reconcile the present ORF3a cryoEM structure with cation conduction. Additional experimental and structural evidence is needed to identify the pathway of cations. The proposed permeation pathway between TM1 and TM2 of both protomers is stabilized by numerous vdW interactions accounting for about -97 kcal/mol. The VDW contribution is the most important compared to the electrostatic energy contribution. This is mainly the result of more than 138 methyl-methyl interactions between the two monomers. Therefore, breaking these interactions to open the potential conduction pore would require a significant amount of energy that should be impossible to reach under a passive condition.

Interestingly, the most prevalent variant of ORF3a given by the analysis of SARS-CoV2 variants is Q57H, which is present in about 25% of sequenced viruses [1]. In the same study, Kern et al. observed that this mutant, located in the TM1 transmembrane segment, does not carry any change in the expression, stability, conductance, selectivity, or gating or the ORF3a channel. These results provide further suggestive evidence that the potential permeation pathway within the core of a dimeric complex might not be the actual conduction route for cations.

In conclusion, we present a modeling study of ORF3a dimeric and tetrameric complex, based on the published CryoEM structure. The results bring more questions than answers. What is the functional relevance of anions populating the central polar cavity? Is the cytosolic domain necessary for conduction? Which route cations take to cross the membrane, and what is the role of Orf3a in transport? And finally, what is the functional oligomeric form of ORF3a? We present these early observations to quickly share these observations if this is useful to guide future experiments. We will publish a more detailed analysis in future communications.

## Acknowledgments

MH was supported by the Intramural Research Program of the NIH (NINDS). This project has been funded in part with federal funds from the National Cancer Institute, National Institutes of Health, under contract HHSN26120080001E. The content of this publication does not necessarily reflect the views or policies of the Department of Health and Human Services, nor does the mention of trade names, commercial products, or organizations imply endorsement by the U.S. Government. FDGN acknowledges the FONDECYT grants 1170733. FDGN and VMM are supported by the Interdisciplinary Center of Neuroscience of Valparaíso, Faculty of Sciences, University of Valparaíso, which is a Millennium Institute supported by the ICM-ANID, project P09-022-F, CINV.

## Methods

A tetrameric model of ORF3a protein (Code: 6XDC) was obtained by overlapping two dimer structures into the cryoEM map deposited in The Electron Microscopy Data Bank (EMDB) (Code: 22138) [1]. A disulfide bond was included to bind the CYS 133 residues of both dimers. The tetrameric structure was protonated according to the standard protonation states at pH 7 and embedded into a POPC (1-palmitoyl-2-oleoyl-phosphatidylcholine) membrane, composed of 544 lipids, and surrounded by 43283 water molecules and ions (122 K+, 122 Cl−), resulting in a salt concentration of 0.5 M. The final dimensions of the system are 166 x 164 x 106 Å3. The system was equilibrated using the six steps scripts provided by the CHARMM-GUI [17–19] webserver under the AMBER molecular package [20–22]. Five hundred ns of molecular dynamics simulations were completed. For proteins, the Amber14sb force field was used [23], LIPID 17 force field for lipids [24], the TIP3P water model [25], and ion parameters reported by Joung and Cheatham [26]. An integration time step of 2 fs was used. The cut-off used for van der Waals interactions was 1.0 nm, and the dispersion correction for energy and pressure was applied. The particle mesh Ewald (PME) method was used [27] to treat electrostatic interactions. The velocity rescale (v-rescale) thermostat [28] was used to keep the temperature constant at 310 K. The semi-isotropic Berendsen barostat88 was used to keep the pressure at 1 atm, 310 K, and 1 bar. MMPBSA.py [29] script was used to calculate MM-GBSA [30] energy values among two monomers. Pore radius profiles were obtained using HOLE software [31] and the MDAnalysis package [32]. We used Python 3 [33] and the packages NumPy [34], pandas [35], and matplotlib [36] for data analysis. Visual Molecular Dynamics software was used for visualization and to generate images [10]. Electrostatic potential maps were computed using APBS [12] and PDB2PQR to obtain the input files [37].

## Notes

### Competing Interest Statement

The authors have declared no competing interest.

