## Supplemental Information for "Analysis of SARS-CoV-2 ORF3a structure reveals chloride binding sites"

**Figures Caption**

Figure S1. Density of the molecular system chemical species, obtained as an average over the last 100 ns of MD. Lipid heads are depicted in green, chloride ions in blue, and potassium ions density in red. Chloride ions profile evidence a peak close to z = -10 Å, while potassium ions accumulate with a higher propensity close to the intracellular side (z < -10).

**Figure S1**

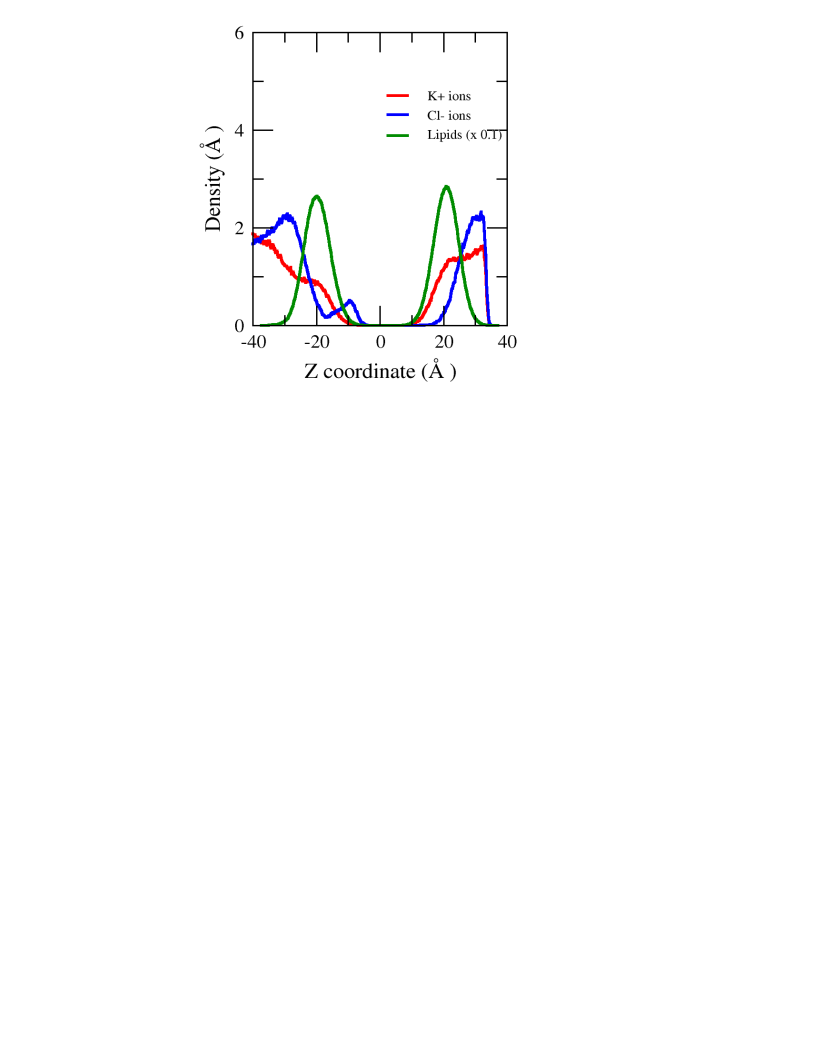
